# Honey varietals differentially impact *Bifidobacterium animalis* ssp *lactis* survivability in yogurt through simulated *in vitro* digestion

**DOI:** 10.1101/2023.10.23.563627

**Authors:** David A. Alvarado, Luis Alberto Ibarra-Sánchez, Annemarie R. Mysonhimer, Tauseef A. Khan, Rong Cao, Michael J. Miller, Hannah D. Holscher

**Author notes:** **Corresponding Author**: Hannah Holscher, 260 Edward R. Madigan Laboratory, 1201 West Gregory Drive, Urbana, IL 61801. Authors’ Last Names: Alvarado, Ibarra-Sánchez, Mysonhimer, Khan, Cao, Miller, Holscher.

## Abstract

**Background:** *Bifidobacterium animalis ssp. lactis* DN-173 010/CNCM I-2494 (*B. animalis*) is a probiotic strain commonly added to yogurt. Yogurt and honey are a popular culinary pairing. Honey improves bifidobacteria survival *in vitro*. However, probiotic survival in yogurt with honey during *in vitro* digestion has not been investigated.

**Objective:** The study aimed to evaluate the effects of different honey varietals and concentrations on *B. animalis* survivability in yogurt through *in vitro* digestion.

**Methods:** Yogurt with honey or control-treated samples underwent *in vitro* simulated oral, gastric, and intestinal digestion. *B. animalis* cells were enumerated on MRS medium followed by an overlay with a modified selective MRS medium; all underwent anaerobic incubation. *B. animalis* were enumerated pre-digestion and after oral, gastric, and intestinal digestion. There were two study phases: phase 1 tested four honey varietals at 20% w/w per 170g yogurt, and phase 2 tested seven dosages of clover honey (20, 14, 10, 9, 8, 6, and 4% w/w) per 170g yogurt.

**Results:** Similar *B. animalis* counts were observed between all treatments after oral and gastric digestion (<1 Log CFU/g probiotic reduction). Higher *B. animalis* survivability was observed in yogurt with clover honey after exposure to simulated intestinal fluids (∼3.5 Log CFU/g reduction; *P* < 0.05) compared to all control treatments (∼5.5 Log CFU/g reduction; *P* < 0.05). Yogurt with 10 to 20% w/w clover honey increased *B. animalis* survivability after simulated *in vitro* digestion (up to ∼4.7 Log CFU/g survival; *P* < 0.05).

**Conclusion:** Yogurt with added honey improves probiotic survivability during *in vitro* digestion. The effective dose of clover honey in yogurt was 10 to 20% w/w per serving (1 – 2 tablespoons per 170g yogurt) for increased probiotic survivability during *in vitro* digestion.

## Introduction

Yogurt is a fermented dairy product (1), created from spontaneous or induced lactic acid fermentation of milk (2,3). The microorganisms used to ferment the milk inform their characterization as standard or probiotic yogurts. Conventional yogurts use a standard starter culture (*Lactobacillus delbrueckii* ssp. *bulgaricus* and *Streptococcus thermophilus)* (4). Probiotic yogurts use the required standard culture in addition to supplemention with probiotic strains, typically Bifidobacterial and/or lactobacilli (5).

Probiotics are defined as “live microorganisms that when administered in adequate amounts confer a health benefit on the host” (6). Probiotic microbial strains must be 1) identified genetically (strain-specific), 2) safe for intended use, 3) supported by at least one human clinical trial, 4) demonstrate health benefit(s), and 5) alive in sufficient numbers in the product at an efficacious dose throughout its shelf life (7,8). Although many fermented foods contain live and active cultures, few qualify as probiotic foods as they do not contain microbes that meet the above conditions (9). Other factors that prevent certain fermented dairy products from having a probiotic status are their capacity to survive through high-stress environments such as gastrointestinal digestion (10). An adequate number of viable cells in probiotic yogurts (10^8^ cells per gram) are necessary to provide strain-specific health benefits, allowing greater opportunity to colonize the intestine (10). Many commercial probiotic strains are from the *Bifidobacterium* genus (11). *B. animalis* is a typical inhabitant of the mammalian colon (12). *B. animalis* is resistant to acidity, adheres to intestinal mucin, grows in milk, and demonstrates some oxidative stress resistance (12).

Certain food combinations can optimize nutrient bioavailability (e.g., carrots consumed with oil enhance carotenoid absorption). Also, there is growing evidence that consuming live microbes in the diet helps support health (13). As yogurt (a source of live microbes and probiotics) is commonly paired with honey, and honey can enhance bifidobacterial survival *in vitro* (14–16), this study aimed to evaluate the effect of adding four different honey varietals (alfalfa, buckwheat, clover and orange blossom) to commercial yogurt containing probiotic *Bifidobacterium animalis ssp. lactis* DN-173 010/CNCM I-2494 (*B. animalis*), on the probiotic survivability within yogurt during *in vitro* digestion. We hypothesized that honey would enhance *B. animalis* survival during simulated complete *in vitro* gastrointestinal digestion.

## Materials and Methods

### Honey Characterization

The National Honey Board provided the honey varietals (alfalfa, buckwheat, clover, and orange blossom). Honey from a single production was packaged in 1-pound containers for retail sale in North America was shipped directly from the supplier to our labs and simultaneously to the analytic lab for composition testing. Within 24 h of receipt, samples were stored at −20°C in airtight 1-pound packages, and aliquots for experimentation were stored at −80°C. Products remained frozen until prepared for use within 10 days of removal from the freezer. The producer used general industry practices to yield honey free of foreign organic matter (heated to ≤185°F, filtered to 16 microns, and cooled to 125°F for packaging). Honey was tested by the producer to ensure authenticity. The honey varietals originated from different locations: clover was from the Dakotas, alfalfa from Wyoming, orange blossom from Orange Groves, and buckwheat from the Midwest.

### Sugar analysis of honey varietals

The sugars from the honey varietals (glucose, fructose, and sucrose) were quantified using high performance anion exchange chromatography with pulsed amperometry detection (HPAEC-PAD, Dionex ICS-5000, Thermo-Fisher, USA) in conjunction with CarboPac PA1 guard (4 mm x 50 mm) and analytical (250 mm x 4 mm) column. The sugars were eluted at 25°C in 10 mM NaOH for 15min, followed by 100 mM NaOH for 30 min at a 1 ml/min flow rate. Honey samples were diluted in deionized water and filtered through a 0.45 µm nylon filter prior to injection on the chromatographic system. Calibration curves were constructed from pure standards (Sigma-Aldrich, St. Louis, MO) were used to quantify honey sugars.

### Antioxidative and phenolic analysis of honey varietals

The total phenolic content (TPC) in honey was determined by the Folin-Ciocalteu method as described previously, with minor modifications (17). Briefly, honey samples were diluted to 30% (w/v) solution with distilled water. Twenty-five µL of diluted sample or standard (gallic acid) solution was mixed with 125 µL 0.2 mol/L Folin-Ciocalteu reagent in a 96-well microplate and allowed to react for 10 min at room temperature. Then 125 µL 7.5% (w/v) Na_2_CO_3_ was added and incubated for 60 min at room temperature. The absorbance was measured at 765 nm using a visible–UV microplate kinetic reader (EL 340, Bio-Tek Instruments, Inc., Winooski, VT, USA). TPC was expressed as mg gallic acid equivalents (GAE) per 100 g honey (mg GAE/100g honey) by using the gallic acid calibration curve.

The 2,2-Diphenyl-1-picrylhydrazyl (DPPH) assay was used to assess the antioxidant activity of honey; it was measured according to a previous report (17), with slight modifications. Briefly, 25 µL of 30% (w/v) honey sample or standard (Trolox) was mixed with 200 µL of 350 µM DPPH in methanol in a 96-well plate. The mixtures were reacted for 6 h in darkness at room temperature. The absorbance was measured at 517 nm. The DPPH antioxidant activity was expressed as µmol of Trolox equivalents (TE) per 100 g honey (µmol TE/100g honey) by plotting the percentage of DPPH quenched against the concentration of Trolox.

Ferric-reducing antioxidant power (FRAP) activity of honey was measured following a previously reported procedure (17) with slight modifications. Briefly, 10 µL of 30% (w/v) honey sample or standard (ascorbic acid) was allowed to react with 300 µL of ferric-TPTZ reagent and kept at room temperature for 2 h. The absorbance was read at 593 nm. The FRAP value was expressed as µmol L-ascorbic acid equivalent per 100 g honey (µmol AAE/100g honey) using the L-ascorbic acid calibration curve.

The phenolic extract of honey was prepared using acidified aqueous methanol. Briefly, honey samples were diluted to 30% (w/v) solution by distilled water and were acidified by formic acid with the final concentration of 1% (v/v). Twenty-five mL of acidified honey solution was purified using OASIS HLB polymeric solid phase extraction cartridges (150 mg, Waters, Mississauga, ON, Canada) and eluted with 1% formic acid in methanol (v/v). The eluent was used for LC-MS analysis. LC-MS/MS analysis was performed using a Thermo® Scientific Q-Exactive™ Orbitrap mass spectrometer equipped with a Vanquish™ Flex Binary UPLC System (Waltham, MA, USA). A Kinetex XB-C18 100A column (100 x 4.6 mm, 2.6 µm, Phenomenex Inc., Torrance, CA, USA) was used. The binary mobile phase consisted of solvent A (99.9% H_2_O/ 0.1% formic acid) and solvent B (94.9% MeOH/ 5% ACNI/ 0.1% formic acid). The following solvent gradient was used: 0 – 5 min, 0% to 12% B; 5 – 15 min, 12% to 23% B; 15 – 30 min, 23% to 50% B; 0 - 40 min, 50% to 80% B; 40 – 42 min, 80% to 100% B; 42 – 45 min, 100% B; 45 – 46 min, 100% to 0% B; 46 – 52 min, 0% B. The column temperature was set at 40°C, the flow rate was set at 0.7 mL/min, and the injection volume was 2 µL; UV peaks were monitored at 280 nm. Mass spectrometry data were collected using both FullMS and DDMS2 modes, negative ionization mode was used; spray voltage was set at 4.5 kV. FullMS was used for quantification, and DDMS2 (TopN=10, NCE = 30, intensity threshold = 1.0e5 counts) was used for the tentative identification of the unknown compounds. Data was visualized and analyzed using Thermo FreeStyle™ 1.7PS2 software.

### Organic acid analysis of honey varietals

Honey samples for organic acid analysis were diluted to 1% (w/v) solution by 1% (v/v) formic acid in distilled water. Further dilution of 0.25% (w/v) was prepared to analyze gluconic acid. Samples were filtered by 0.45 µm syringe filters before LC-MS analysis. LC-MS/MS analysis was performed using the same HPLC-MS system as above stated. A Phenomenex Rezex^TM^ ROA-Organic Acid H^+^ (8%) column (150 x 4.6 mm, Phenomenex Inc., Torrance, CA, USA) was used. The mobile phase was 0.5% formic acid in water. Separation was achieved using isocratic elution with a flow rate of 0.3 mL/min, method duration was 7 min. The column temperature was set at 55°C and the injection volume was 0.5 µL. Mass spectrometry data were collected using the FullMS method, negative ionization mode was used and the spray voltage was set at 4.0 kV. Data were visualized using Thermo FreeStyle™ 1.7PS2 software. All analyses were performed in triplicate.

### Enzymatic analysis of honey varietals

Amylase activity in the honey varietals was measured by diluting the honey samples to 30 % (w/v) solution by distilled water. The solution was filtered through a 0.22 µm filter to remove any insoluble materials. Amylase activities were measured using a colorimetric assay kit (Abcam, Waltham, MA, USA) according to the manufacturer’s instructions. Briefly, 50 µL of diluted honey samples or nitrophenol standards were mixed with 100 µL of amylase reaction mix (ethylidene-pNP-G7 and α-glucosidase) in a 96-well plate. Measure absorbance immediately at 405 nm in a kinetic mode for 60 min at 25°C protected from light. α-Amylase in honey cleaved the substrate ethylidene-pNP-G7 to produce smaller fragments that were eventually modified by α-glucosidase, causing the release of a chromophore that can be measured at 405 nm. The amylase activity was expressed as U /100 g honey by using the nitrophenol calibration curve. One U was defined as amount of amylase that cleaves ethylidene-pNP-G7 to generate 1.0 μmol of nitrophenol per min at pH 7.20 at 25 °C.

Diastase activity in the honey varietals was measured by diluting the honey samples to 1 % (w/v) solution by 0.1M acetate buffer (pH = 5.2). The solution was filtered through a 0.22 µm filter to remove any insoluble materials. Diastase activities were measured using a colorimetric assay kit (Phadebas, Cambridge, MA, USA) according to the manufacturer’s instructions. Briefly, 5.0 mL of diluted honey samples or 0.1M acetate buffer using as blank were mixed with one Phadebas® tablet at 40 °C for 30 min. The Phadebas® tablet contained 45 mg of water-insoluble, cross-linked starch polmer carrying blue dye, which can be hydrolyzed by diastase and generate blue water-soluble fragments. The reaction was stopped by adding 1 mL of 0.5M sodium hydroxide solution. After centrifuging at 1500 x g for 5 min, the supernatant were measured in 1 cm cuvette at 620nm. Diastase activity was expressed as diastase number (DN) based on the difference of absorption at 620 nm between sample and blank.

Glucose oxidase activity in the honey varietals was measured by diluting the honey samples to 30 % (w/v) solution by distilled water. The solution was filtered through a 0.22 µm filter to remove any insoluble materials. Glucose oxidase activities were measured using a colorimetric assay kit (Abcam, Waltham, MA, USA) according to the manufacturer’s instructions. Briefly, 50 µL of diluted honey samples or glucose oxidase standards were mixed with 50 µL of glucose oxidase reaction mix (glucose, AbRed indicator, and horseradish peroxidase) in a 96-well plate. Measure absorbance immediately at 570 nm in a kinetic mode for 30 min at 37°C. Glucose oxidase in samples catalyzed the oxidation of β-D-glucose into hydrogen peroxide and D-glucono-1,5-lactone. The produced hydrogen peroxide reacted with AbRed indicator when catalyzed by horseradish peroxidase to generate the compound which can be measured at 570 nm. The glucose oxidase activity was expressed as U /100 g honey by using the calibration curve. One U was defined as amount of glucose oxidase that react with 1.0 μmol of glucose per min at 37 °C.

Catalase activity was measured by diluting the honey samples to 3.0 % (w/v) solution by distilled water. The solution was filtered through a 0.22 µm filter to remove any insoluble materials. Catalase activities were measured using a colorimetric assay kit (Cayman Chemical, Ann Arbor, MI, USA) according to the manufacturer’s instructions. Briefly, 20 µL of diluted honey samples or formaldehyde standards were mixed with 100 µL of assay buffer, 30 µL of methanol, and 20 µL of 35.3 mM hydrogen peroxide in a 96-well plate. Catalase in samples catalyzed the peroxidation of methanol to produce formaldehyde after 20 min incubation at room temperature. The formaldehyde was measured calorimetrically at 540 nm with 4-amino-3-hydrazino-5-mercapto-1,2,4-triazole (purpuld) and potassium periodate. The catalase activity was expressed as U /100 g honey by using the formaldehyde calibration curve. One U was defined as amount of catalase that peroxidate methanol to generate 1.0 μmol of formaldehyde per min at room temperature.

Invertase activity was measured by diluting the honey samples to 30 % (w/v) solution with distilled water. The solution was filtered through a 0.22 µm filter to remove any insoluble materials. Glucose in honey was removed by centrifuging at 5,000 x g using an Amicon ultra centrifugal filter device with a molecular weight cut-off of 10,000 (Merck KGaA, Darmstadt, Germany) for at least seven times. The concentrated protein was collected and dissolved up to 500 µL in PBS buffer. Invertase activities were measured using a colorimetric assay kit (Abcam, Waltham, MA, USA) according to the manufacturer’s instructions. Briefly, 25 µL of concentrated protein solution was mixed with 15 µL of assay buffer and 10 µL of sucrose (i.e. invertase substrate). The same sample volume without adding sucrose was prepared simultaneously as a background control. Invertase in honey catalyzed the hydrolysis of sucrose by cleaving its glycosidic bond and forming glucose and fructose. After 20 min of reaction, samples, background controls, and sucrose standards were mixed with the provided enzyme mix and probe to generate a chromogen that can be measured at 570 nm. The absorption of background control was subtracted from the sample to eliminate the influence of residual glucose in a sample. The invertase activity was expressed as mU /100 g honey by using the glucose calibration curve. One mU was defined as amount of invertase that cleaves sucrose to generate 1.0 nmol of glucose per min at 37 °C.

### In Vitro Experimentation Phase 1 (Comparison of Honey Varietals)

Honey varietals were stored at −80°C until the experiment began and then thawed in a water bath at 42°C for 30 minutes until a syrup consistency was observed. The yogurt used in this study was a commercial low-fat vanilla yogurt containing *B. animalis* (according to the manufacturer’s label) throughout the experiment. For the first phase of the in vitro experimentation, probiotic-containing samples were prepared by adding the four honey varietals to each yogurt sample to a final concentration of approximately 20% w/w in the yogurt. Each sample contained 170 g of commercial yogurt plus 42 g of honey or a control component. The controls for the first experiment were an undiluted yogurt, yogurt with added water (20% w/v), and 30.4 g sucrose (isocaloric equivalent to the 42g honey). After treatment preparation, all samples were stored at 4°C for 72 hours to allow *B. animalis* to acclimate to its new yogurt matrix.

Following Brodkorb’s protocol with scaled-down modifications (16), enzymatic assays (α-salivary amylase, porcine pepsin, pancreatin, trypsin, and bile salt) were conducted 24 h before the experiment, and activity values were considered valid for one week. For the oral stage of digestion, a spectrophotometric stop reaction was used to calculate the activity of human salivary α-amylase (Sigma-Aldrich, cat. No. 1031) using Equation 1 (b = intercept of linear regression, a = slope of linear regression, X = quantity of amylase powder (mg) added before stopping the reaction): Units/mg = ([A_540_ Test − A_540_ Blank] – b) / (a × X). Each assay replicate had final amylase activity of 75 U/mL using equation 1. The gastric stage of digestion involved assessing pepsin activity using a spectrophotometric stop reaction (16). First, pepsin was measured to 12,000 U, then diluted to a final concentration of 2,000 U/mL using deionized water. Next, 360 U gastric lipase was diluted to 60 U/mL using deionized water (16). Finally, the activity was calculated using Equation 2 (X = mg quantity of pepsin powder): Units/mg = ([A_260_ Test – A_260_ Blank] × 1000) / (Δt × X × 0.001). For the intestinal stage of digestion, we conducted two assays to measure the trypsin activity in pancreatin using the Brodkorb et al. protocol with modifications (16). First, 800 U pancreatin was diluted to a final activity of 100 U/mL using deionized water. Next, bile salts were supplied from bovine bile and measured using a commercial kit according to the supplier’s protocol (Sigma-Aldrich, cat. No. MAK 309). Finally, trypsin in pancreatin was measured using a kinetic spectrophotometric rate determination method and calculated using Equation 3 (X = quantity of pancreatin used in the final reaction mixture in mg): Units/mg = ([A_260_ Test – A_260_ Blank] × 1000 × 3) / (540 × X).

All yogurt samples underwent *in vitro* simulation of gastrointestinal digestion using the Brodkorb et al. protocol with modifications (16). For each stage [oral, gastric, and intestinal], a simulated digestive fluid was prepared and labeled as simulated salivary, gastric, and intestinal fluids, respectively. Each solution was prepared 72 h prior to starting a trial for the experiment and stored at 4°C. For each trial, an aliquot was pre-warmed to 37°C the same day of the experiment (16). The electrolyte solutions used for each were prepared using the following quantities (deionized water was used in all dilutions): 0.5M KCl, 0.5M KH_2_PO_4_, 1M NaHCO_3_, 2M NaCl, 0.15M MgCl_2_(H_2_O)_6_, 0.5M (NH_4_) _2_CO_3_, 6M HCl (as needed to achieve required pH), 0.3M CaCl_2_(H_2_O) _2_ (16). Each simulated fluid (salivary, gastric, and intestinal) was prepared by mixing the following volumes of electrolyte solutions and diluting with water to a final volume of 400 mL to achieve the indicated final mM concentrations, and adjusted to pH 7, 3 and 7 respectively. Simulated salivary solution had a pH of 7 and contained 15.1 mM KCl (15.1 mL), 3.7 mM KH_2_PO_4_ (3.7 mL), 13.6 mM NaHCO_3_ (6.8 mL), 0.15 mM MgCl_2_(H_2_O)_6_ (0.5 mL), 0.06 mM (NH_4_)_2_CO_3_ (0.06 mL), 1.1 mM HCl (0.09 mL), 1.5 mM CaCl_2_(H_2_O)_2_ (0.025 mL). The simulated gastric fluid had a pH of 3, and contained 6.9 mM KCl (6.9 mL), 0.9 mM KH_2_PO_4_ (0.9 mL), 25 mM NaHCO_3_ (12.5 mL), 47.2 mM NaCl (11.8 mL), 0.12 mM MgCl_2_(H2O)_6_ (0.4 mL), 0.5 mM (NH_4_)_2_CO_3_ (0.5 mL), 15.6 mM HCl (1.3 mL), 0.15 mM CaCl_2_(H_2_O)_2_ (0.005 mL) (16). The simulated intestinal fluid had a pH of 7 and contained 6.8 mM KCl (6.8 mL), 0.8 mM KH_2_PO_4_ (0.8 mL), 42.5 mM NaHCO_3_ (85 mL), 9.6 mM NaCl (38.4 mL), 0.33 mM MgCl_2_(H2O)6 (1.1 mL), 8.4 mM HCl (0.7 mL), 0.6 mM CaCl_2_(H_2_O)_2_ (0.04 mL) (16).

The sequential simulated digestion procedure was conducted, as indicated in **Table 1**, by mixing the sample with the appropriate solutions in a sterile 15 mL polypropylene centrifuge tube followed by incubation at 37°C (Table 1). For all experiments, at four-time points—pre-digestion (baseline) and after each stage of digestion (i.e., oral, gastric, and intestinal)—2 mL aliquots were removed and placed in ice to slow and stop the enzymatic activity. Sample aliquots were serially diluted 10-fold using a phosphate-buffered saline (PBS) solution. Enumeration was carried out by spread plating 100 µL of each dilution factor onto separate ∼20 mL of solidified MRS agar supplemented with 0.5 g/L L-cysteine hydrochloride (MRSc) and then incubated for 5 h at 37°C under anaerobic conditions to allow *B. animalis* cells to recover (18). Plates were then overlaid with a selective media (∼20 mL): MRSc supplemented with lithium chloride (3 g/L) and sodium propionate (4.5 g/L) and incubated at 37°C for an additional 67 h under anaerobic conditions before the enumeration of *B. animalis* (19). The incubation time was sufficient to obtain visible colonies on the plates displaying typical *Bifidobacterium* morphology (18). Additionally, we tested a heat-treated yogurt with the selective medium to ensure that no detectable amounts of bifidobacteria would grow after pasteurization; as soon as the internal temperature of the yogurt reached 63 C and then started a 30 min timer.

**Table 1.** Sequential simulated *in vitro* digestion protocol.

|  |
| --- |
| **Oral stage (final volume 5mL, pH 7)** |
| Food-oral fluid ratio 1:1 (vol/vol) |
| 2.5g yogurt sample |
| 2mL SSF electrolyte (prewarmed 30 min at 37°C) |
| 12.5 µL CaCl<sub>2</sub>(H<sub>2</sub>O)<sub>2</sub> (final concentration 1.5 mM) |
| 112.5 µL H<sub>2</sub>O |
| 375 µL salivary amylase (1,000 U/mL) – 375 U – Final 75 U/mL |
| 30 min incubation at 37°C |
| **Gastric stage (final volume 6mL, pH 3)** |
| Oral-gastric fluid ratio 1:1 (vol/vol) |
| 3mL oral stage |
| 2.4mL SGF electrolyte (prewarmed 30 min at 37°C) |
| 1.5 µL CaCl<sub>2</sub>(H<sub>2</sub>O)<sub>2</sub> (final concentration 0.15 mM) |
| 362.5 µL H<sub>2</sub>O |
| 36 µL HCl (6M) |
| 0.2 mL porcine pepsin (60,000 U/mL) – 12,000 U – Final 2000 U/mL |
| 120 min incubation at 37°C |
| **Intestinal stage (final volume 8mL, pH 7)** |
| Gastric-intestinal fluid ratio 1:1 |
| 4mL gastric stage |
| 1.6mL SIF electrolyte (prewarmed 30 min at 37°C) |
| 748 µL H<sub>2</sub>O |
| 52 µL NaOH (5M) |
| 0.6 mL Bile salts (Final 10 mM) (prewarmed 30 min at 37°C) |
| 1mL pancreatin (800 U/mL trypsin) – 800 U – Final 100 U/mL |
| 120 min incubation at 37°C |

All experiments were run independently three times with triplicate samples for each time point, and dilutions were plated in triplicate to obtain an average for each trial.

### In Vitro Experimentation Phase 2 (Dose Response)

Based on results from Phase 1, which compared the effect of four different honey varietals on probiotic survivability, we identified clover honey as having the greatest effect on supporting *B. animalis* survivability *in vitro* compared to the other three honey varietals. Thus, we conducted a second phase of experimentation to assess a dose relationship for clover honey. The dosages used in this second experiment were: 0 g (0% w/w), 7 g (4% w/w), 10.5 g (6% w/w), 14 g (8% w/w), 17.5 g (9% w/w), 21 g (10% w/w), 28 g (14% w/w), 42 g (20% w/w) of honey added to 170 g of yogurt. All samples followed the same procedure as outlined in Phase 1. All experiments were run independently three times with triplicate samples for each time point, and dilutions were plated in triplicate to obtain an average for each trial.

### Statistical analyses

Honey characterization data was visualized and analyzed using Thermo FreeStyle™ 1.7PS2 software. *In vitro* digestion experiments were analyzed as completely randomized designs using JMP 13.1 (SAS Institute Inc., Cary, NC). An analysis of variance (ANOVA) was performed to establish the significance of factor (honey treatments). Honey characterization results (sugar, phenolic, enzymes, and organic acid profiles) and probiotic enumeration results (log CFU/g) were evaluated using Tukey’s test to determine the statistical significance of mean differences between treatments. Additionally, for probiotic enumeration results, a Dunnett test was used to compare the honey treatment dosages to the control (undiluted yogurt).

## Results

For each honey varietal, the concentrations of glucose, fructose, and sucrose were determined as a proportion of the honey (g/100g varietal), **Table 2**. Fructose did not differ between the varietals; however, there were differences in sucrose and glucose concentrations among the varietals, with clover honey having the greatest amounts of both sugars compared to the other varietals. The phenolic composition of the honey varietals is reported in **Table 2**. The clover honey had 10%, 53%, and 36% higher concentrations of *trans*-ferulic acid and 14%, 78%, and 34% higher concentrations of kaempferol compared to the buckwheat, orange, and alfalfa varietals, respectively. Among the enzymes assessed, amylase, diastase, glucose oxidase, and invertase differed among the varietals, **Table 2**. Clover honey had the lowest amounts of both glucose oxidase and invertase among the varietals.

**Table 2.** Honey composition of the four varietals at baseline.

| | Honey varietals (mean ± SEM) | | | |
| --- | --- | --- | --- | --- |
|  | Clover | Buckwheat | Orange | Alfalfa |
| **Sugars (g/100g honey)** |  |  |  |  |
| Fructose | 38.5 ± 0.30 | 37.5 ± 1.01 | 38.2 ± 0.25 | 36.6 ± 0.83 |
| Glucose | 34.6 ± 0.11<sup>a</sup> | 33.6 ± 0.68<sup>ab</sup> | 31.8 ± 0.21<sup>b</sup> | 32.5 ± 0.50<sup>b</sup> |
| Sucrose | 0.72 ± 0.01<sup>a</sup> | 0.12 ± 0.01<sup>b</sup> | 0.68 ± 0.01<sup>a</sup> | 0.42 ± 0.01<sup>c</sup> |
| **Phenolic (μg/100g honey)** |  |  |  |  |
| *p*-Hydroxybenzoic Acid | 391 ± 18<sup>b</sup> | 763 ± 28<sup>a</sup> | 354 ± 8.0<sup>b</sup> | 240 ± 13<sup>c</sup> |
| Caffeic Acid | 130 ± 6.0<sup>c</sup> | 190 ± 11<sup>c</sup> | 1300 ± 41<sup>a</sup> | 1200 ± 38<sup>b</sup> |
| *p*-Coumaric Acid | 330 ± 14<sup>b</sup> | 530 ± 1.9<sup>a</sup> | 160 ± 8.0<sup>c</sup> | 180 ± 8.0<sup>c</sup> |
| Trans-Ferulic Acid | 150 ± 5.0<sup>a</sup> | 140 ± 2.0<sup>b</sup> | 70. ± 2.0<sup>d</sup> | 97 ± 3.0<sup>c</sup> |
| Abscisic Acid | 268 ± 10.<sup>b</sup> | 244 ± 9<sup>b</sup> | 1070 ± 4<sup>a</sup> | 170. ± 9<sup>c</sup> |
| Kaempferol-rutinoside | 13 ± 1.0<sup>c</sup> | 88 ± 3.0<sup>a</sup> | 15 ± 1.0<sup>c</sup> | 39 ± 2.0<sup>b</sup> |
| Pinobanksin | 1100 ± 15<sup>c</sup> | 1300 ± 46<sup>b</sup> | 420 ± 14<sup>d</sup> | 1500 ± 3<sup>a</sup> |
| Naringenin | 490 ± 8.0<sup>c</sup> | 630 ± 22<sup>b</sup> | 150 ± 7.0<sup>d</sup> | 698 ± 15<sup>a</sup> |
| Quercetin | 93 ± 2.0<sup>b</sup> | 77 ± 4.0<sup>c</sup> | 110 ± 2.0<sup>a</sup> | 75 ± 3.0<sup>c</sup> |
| Kaempferol | 690 ± 22<sup>a</sup> | 598 ± 27<sup>b</sup> | 150 ± 4.0<sup>d</sup> | 460 ± 6.0<sup>c</sup> |
| Apigenin | 280 ± 6.0<sup>b</sup> | 290 ± 12<sup>b</sup> | 72 ± 1.0<sup>c</sup> | 360 ± 9.0<sup>a</sup> |
| Pinocembrin | 1600 ± 34<sup>a</sup> | 1500 ± 51<sup>a</sup> | 240 ± 4.0<sup>b</sup> | 1600 ± 41<sup>a</sup> |
| Biochanin A | 320 ± 12<sup>b</sup> | 420 ± 18<sup>a</sup> | 92 ± 2.0<sup>c</sup> | 410 ± 13<sup>a</sup> |
| Chrysin | 270 ± 10.<sup>c</sup> | 370 ± 12<sup>b</sup> | 72 ± 2.0<sup>d</sup> | 510 ± 17<sup>a</sup> |
| Galangin | 290 ± 8.0<sup>b</sup> | 380 ± 20.<sup>a</sup> | 75 ± 3.0<sup>c</sup> | 301 ± 11<sup>b</sup> |
| **Organic acids (mg/100g honey)** |  |  |  |  |
| Citric Acid | 5.7 ± 0.11<sup>c</sup> | 6.10 ± 0.10<sup>b</sup> | 11 ± 0.07<sup>a</sup> | 5.3 ± 0.02<sup>d</sup> |
| Gluconic Acid | 370 ± 3.3<sup>c</sup> | 430 ± 5.90<sup>b</sup> | 510 ± 1.8<sup>a</sup> | 503 ± 1.2<sup>a</sup> |
| Malic Acid | 2.8 ± 0.04<sup>d</sup> | 5.20 ± 0.07<sup>b</sup> | 6.40 ± 0.13<sup>a</sup> | 3.5 ± 0.03<sup>c</sup> |
| Succinic Acid | 0.90 ± 0.04<sup>c</sup> | 1.20 ± 0.0<sup>b</sup> | 1.60 ± 0.03<sup>a</sup> | 1.1 ± 0.09<sup>b,c</sup> |
| Lactic Acid | 2.0 ± 0.07<sup>d</sup> | 3.20 ± 0.09<sup>c</sup> | 6.50 ± 0.08<sup>a</sup> | 5.2 ± 0.14<sup>b</sup> |
| **Total phenolic activity** |  |  |  |  |
| TPC (GAE mg/100g) | 36 ± 0.50<sup>d</sup> | 195 ± 2.0<sup>a</sup> | 48 ± 0.50<sup>b</sup> | 43 ± 0.50<sup>c</sup> |
| FRAP (AAE μmol/100 g) | 127 ± 2.1<sup>c</sup> | 571 ± 5.9<sup>a</sup> | 146 ± 1.1<sup>b</sup> | 152 ± 2.0<sup>b</sup> |
| DPPH (TE μmol/100 g) | 83 ± 4.2<sup>c</sup> | 570 ± 18<sup>a</sup> | 142 ± 6.6<sup>b</sup> | 122 ± 5.8<sup>b</sup> |
| **Enzymes** |  |  |  |  |
| Amylase (U/100g) | 22.5 ± 0.58<sup>b</sup> | 19.4 ± 0.81<sup>c</sup> | 17.3 ± 0.61<sup>d</sup> | 27.9 ± 0.84<sup>a</sup> |
| Diastase (DN) | 7.41 ± 0.10<sup>b</sup> | 5.66 ± 0.09<sup>d</sup> | 6.09 ± 0.07<sup>c</sup> | 8.57 ± 0.06<sup>a</sup> |
| Glucose Oxidase (U/100g) | 0.72 ± 0.01<sup>c</sup> | 4.86 ± 0.01<sup>a</sup> | 1.04 ± 0.02<sup>b</sup> | 0.47 ± 0.02<sup>d</sup> |
| Catalase (U/100g) | 42.7 ± 1.29<sup>ab</sup> | 35.8 ± 1.77<sup>d</sup> | 46.3 ± 2.45<sup>a</sup> | 40.9 ± 1.02<sup>bc</sup> |
| Invertase (mU/100g) | 12.8 ± 0.14<sup>c</sup> | 30.5 ± 4.99<sup>b</sup> | 39.3 ± 0.75<sup>a</sup> | 27.8 ± 0.86<sup>b</sup> |

<sup>a, b, c, d</sup> Means with dissimilar letters in a column are significantly different using Tukey's test ( $P < 0.05$ )
Abbreviations: gallic acid equivalents (GAE); Trolox equivalents (TE); L-ascorbic acid equivalent (AAE)

In the Phase 1 experimentation, each of the four honey varietals (alfalfa, buckwheat, clover, and orange blossom) was compared against three controls (i.e., undiluted, sucrose, and water). The addition of honey did not affect the survial of *B. animalis* in yogurt during the baseline, oral, or gastric phases. During the intestinal phase, clover honey enhanced the survivability of *B. animalis* in yogurt. (**Table 3**; **Figure 1**). After intestinal digestion, clover honey demonstrated the least log CFU/g reduction of *B. animalis* from baseline after complete simulated digestion from baseline (∼3.8 Log CFU/g reduction). This reduction in clover was significantly less compared to the controls: sucrose (∼5.8 Log CFU/g reduction, P < 0.05), water vehicle (∼5.9 Log CFU/g reduction, P < 0.05), and undiluted (∼5.4 Log CFU/g reduction, P < 0.05). After intestinal digestion, alfalfa (∼4.5 Log CFU/g reduction), buckwheat (∼5.5 Log CFU/g reduction), and orange (∼5.6 Log CFU/g reduction) had similar reductions in *B. animalis* counts compared to controls, while clover honey resulted in higher *B. animalis* counts after exposure to simulated intestinal fluids (∼3.8 Log CFU/g reduction) compared to undiluted yogurt, sucrose-added yogurt, and water-added yogurt (∼5.5 Log CFU/g reduction, P ˂ 0.05).

**Figure 1.**
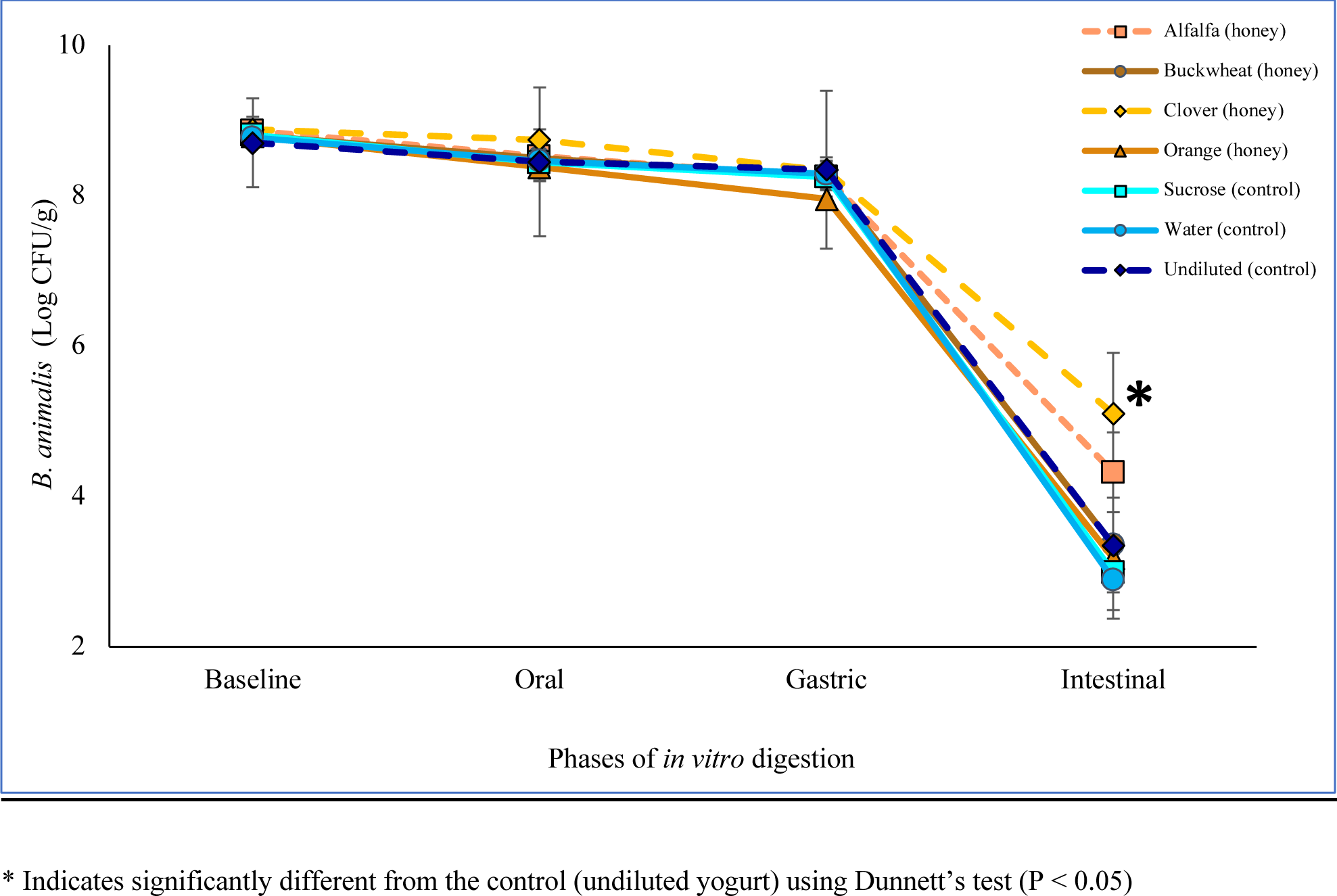
Effect of honey varietals (20% w/w) and controls on *B. animalis* survivability in yogurt (170g) through simulated *in vitro* digestion.

**Table 3.** Effect of different honey varietals on *B. animalis* survivability in yogurt through simulated *in vitro* digestion.

| Yogurt treatment | *B. animalis* (log CFU/g) ± SEM | | | |
| --- | --- | --- | --- | --- |
|  | Baseline:<br>Pre-digestion | 1st stage:<br>Oral | 2nd stage:<br>Gastric | 3rd stage:<br>Intestinal |
| Sucrose | 8.8 ± 0.17 | 8.4 ± 0.25 | 8.3 ± 0.26 | 3.0 ± 0.50<sup>A</sup> |
| Water | 8.8 ± 0.11 | 8.5 ± 0.24 | 8.3 ± 0.12 | 2.9 ± 0.52<sup>A</sup> |
| Undiluted | 8.7 ± 0.59 | 8.5 ± 0.99 | 8.4 ± 1.05 | 3.4 ± 0.12<sup>AB</sup> |
| Orange honey | 8.8 ± 0.08 | 8.4 ± 0.18 | 8.0 ± 0.13 | 3.2 ± 0.10<sup>A</sup> |
| Alfalfa honey | 8.9 ± 0.13 | 8.5 ± 0.23 | 8.3 ± 0.08 | 4.3 ± 0.53<sup>AB</sup> |
| Buckwheat honey | 8.8 ± 0.09 | 8.5 ± 0.18 | 8.3 ± 0.19 | 3.4 ± 0.63<sup>AB</sup> |
| Clover honey | 8.9 ± 0.17 | 8.7 ± 0.14 | 8.3 ± 0.13 | 5.1 ± 0.81<sup>B</sup> |

<sup>A, B</sup> Means with dissimilar letters in a column are significantly different using Tukey's test ( $P < 0.05$ ) to compare all treatment means to the mean of every other treatment. Means with no letters in a column are not significantly different from each other ( $P > 0.05$ ) SEM = Standard Error of the Mean

Based on the improved survivability of the clover honey after intestinal digestion, further experimentation (Phase 2) was initiated to determine a dose-response relationship between clover honey and *B. animalis* survivability (**Table 4**; **Figure 2**). The result revealed a similar reduction to that by the same dose of clover honey in the Phase 1 study, from baseline (∼4.0 Log CFU/g reduction) at the highest dosage and was again significantly different from the undiluted control (∼5.6 Log CFU/g reduction, P < 0.05). Furthermore, lower dosages at 28 g (14% w/w, ∼4.6 Log CFU/g reduction) and 21 g (10% w/w, ∼4.8 Log CFU/g reduction), were also different from the undiluted control (P < 0.05). This indicated a concentration threshold for clover honey that could benefit *B. animalis* survival. Dosages less than 21 g per 170 g yogurt (10% w/w) did not improve *B. animalis* survivability after *in vitro* intestinal digestion and were not different from the control.

**Figure 2.**
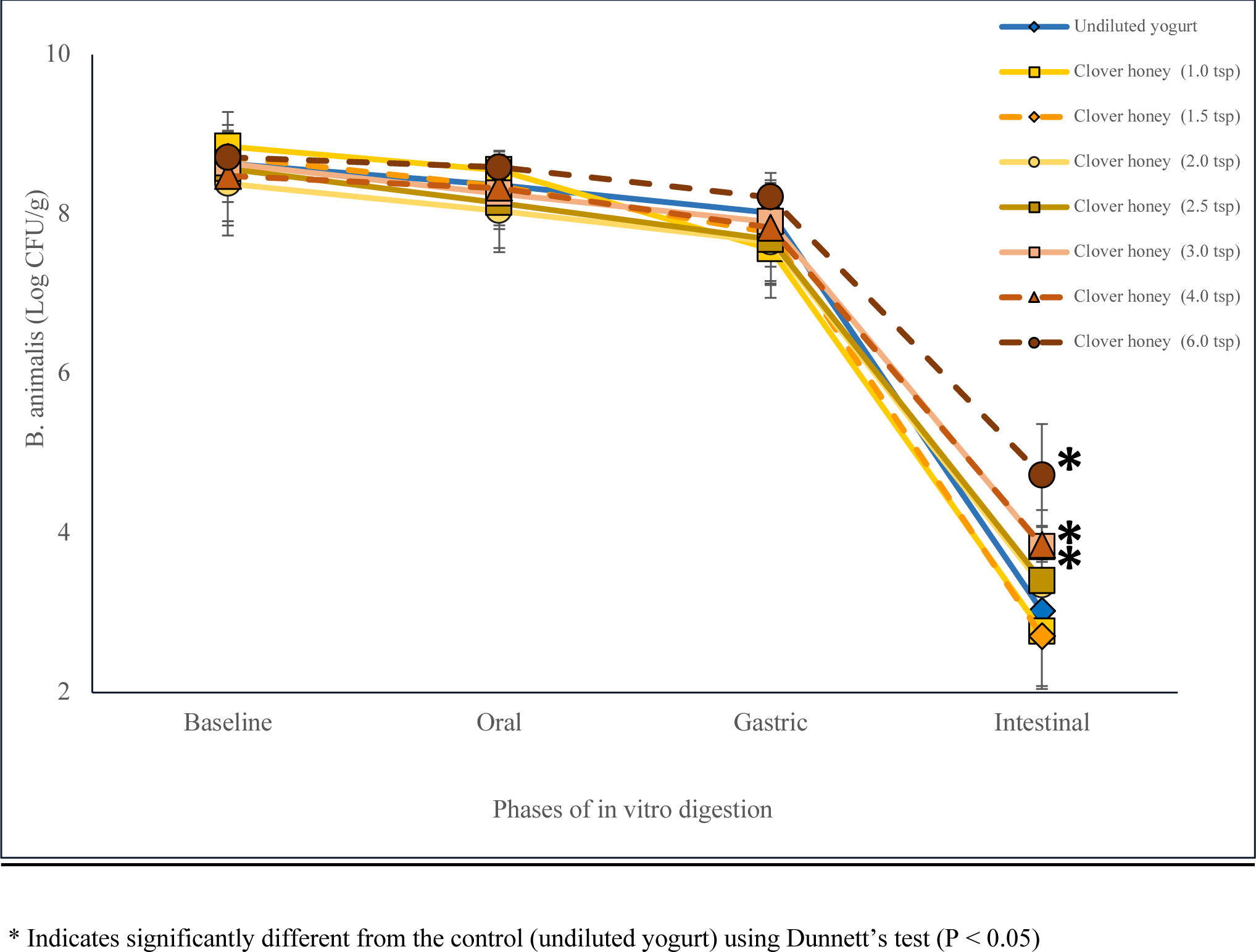
Effect of clover honey at different dosages on *B. animalis* survivability in yogurt (170g) through simulated in vitro digestion.

**Table 4.** Effect of clover honey at different dosages on *B. animalis* survivability in yogurt through simulated *in vitro* digestion.

| Treatment (g of honey<br>per 170g serving of<br>commercial yogurt) | *B. animalis* (log CFU/g) ± SEM | | | |
| --- | --- | --- | --- | --- |
|  | Baseline: pre-digestion | 1st stage:<br>Oral | 2nd stage:<br>Gastric | 3rd stage:<br>Intestinal |
| Undiluted yogurt (0 g) | 8.6 ± 0.38 | 8.4 ± 0.35 | 8.0 ± 0.38 | 3.0 ± 0.37<sup>A, a</sup> |
| Clover honey (7 g) | 8.9 ± 0.19 | 8.6 ± 0.19 | 7.6 ± 0.61 | 2.8 ± 0.72<sup>A, a</sup> |
| Clover honey (10.5 g) | 8.7 ± 0.21 | 8.4 ± 0.36 | 7.8 ± 0.66 | 2.7 ± 0.63<sup>A, a</sup> |
| Clover honey (14 g) | 8.4 ± 0.65 | 8.0 ± 0.52 | 7.7 ± 0.52 | 3.3 ± 0.29<sup>AB, a</sup> |
| Clover honey (17.5 g) | 8.6 ± 0.71 | 8.1 ± 0.57 | 7.7 ± 0.52 | 3.4 ± 0.27<sup>AB, a</sup> |
| Clover honey (21 g) | 8.6 ± 0.48 | 8.3 ± 0.44 | 7.9 ± 0.34 | 3.8 ± 0.26<sup>AB, b</sup> |
| Clover honey (28 g) | 8.5 ± 0.57 | 8.3 ± 0.47 | 7.8 ± 0.49 | 3.9 ± 0.44<sup>AB, b</sup> |
| Clover honey (42 g) | 8.7 ± 0.21 | 8.6 ± 0.19 | 8.2 ± 0.31 | 4.7 ± 0.65<sup>B, b</sup> |

<sup>A, B</sup> Means with dissimilar letters in a column are significantly different using Tukey's test ( $P < 0.05$ ) to compare all treatment means to the mean of every other treatment.
<sup>a, b</sup> Means with dissimilar letters in a column are significantly different from the control (undiluted yogurt) using Dunnett's test ( $P < 0.05$ )

## Discussion

Herein, we evaluated the effect of pairing four honey varietals (alfalfa, buckwheat, clover, and orange blossom) with commercial yogurt on the survivability of *Bifidobacterium animalis ssp. lactis* DN-173 010/CNCM I-2494 through *in vitro* digestion. Our results demonstrated that clover honey supported *B. animalis* survival best against the harsh environments of the simulated *in vitro* intestinal stages of digestion compared to the other honey varietals and the controls. We also observed that the minimum effective dosage of clover honey that improved probiotic survivability after *in vitro* intestinal digestion was 10% w/w in yogurt. Analysis of the nutrient profiles of honey revealed that clover honey had higher concentrations of glucose and sucrose, as well as phenolic compounds kaempferol and trans-ferulic acid compared to other honey varietals. Conversely, components that were low in clover honey (i.e., invertase and glucose oxidase) may also have contributed to the enhanced survivability of *B. animalis*.

*Bifidobacterium* species can utilize sugars with different degrees of polymerization (mono-, oligo-, and polysaccharides) as an energy source (20). The ability to break down various sugar sources is an evolutionary trait stemming from competing for limited carbohydrate sources in the gastrointestinal tract (20). Honey is a natural syrup that is, on average, made up of fructose (39%), glucose (31%), maltose (7%), a variety of oligosaccharides (4%), and sucrose (2%) (21). Honey’s saccharide content, acidity, crop year, production area, granulating tendency, color, plant source, and storage conditions vary depending on its origin. These variables contribute to a unique chemical profile for each honey varietal (22). In addition, the oligosaccharide composition and concentration in honey varietals differ depending on the floral source and honeybee species as the nectar goes is digested by the bee via α-D-glucosidase after collection from flowers (21). Ultimately, the oligosaccharides in honey can be energy substrates for *Bifidobacterium* (14)(1).

Honey varietals are composed of sugars with different concentrations of fructose, glucose, and sucrose. Our study revealed that all four honey varietals varied in fructose and glucose concentrations. Interestingly, enzymatic activity in the honey also differed, with a notably lower invertase in clover compared to the other varietals. Enzymes such as catalase, diastase, and invertase can liberate glucose and fructose from oligosaccharides and disaccharides found within honey (24). Honey ripening has been correlated with the invertase activity and sucrose concentration, and low levels of invertase activity indicate honey quality (25). Glucose oxidase also has the capacity to convert glucose into hydrogen peroxide, an enzyme that was significantly lower in both clover and orange, a compound that is associated with antimicrobial function (24). Our results revealed that clover honey enhanced probiotic survivability through an *in vitro* digestion system compared to the other honey varietals (alfalfa, buckwheat, and orange blossom) and controls. The nutritional analysis revealed that clover honey contained lower concentrations of organic acids (gluconic, malic, and succinic acids) than the other honey varietals. The clover honey also contained greater concentrations of phenolics (trans-ferulic acid and kaempferol) compared to the other honey varietals. These viable antioxidative compounds and reducing sugars may have been an active component only when using 10% clover honey or greater in 170 g of yogurt, as anything less may not have been sufficient to protect *B. animalis*. Other studies have demonstrated that *Bifidobacterium* and LAB strains can transform ferulic acid into p-coumaric acid and caffeic acid, potentially creating a favorable environment for themselves (26). Another study using a LAB strain demonstrated that overexposure to p-coumaric acid induced a stress-induced adaptive response that increased cell surface membrane proteins, to counteract the phenolic toxic levels, similarly observed in the gut environment (27). Follow-up studies are needed to assess if the probiotic in this study may have also gone through a similar adaptive response during the 72 h storage period before the *in vitro* digestion process, and more specifically, what gene or function was induced as a result of a similar adaptation described by Reveron et al.

Herein, clover had significantly less gluconic acid compared to the other varietals. Interestingly, gluconic acid can be utilized by most bifidobacteria strains except for the *B. animalis* species (Asano et al., 1994). Thus, the higher concentration of gluconic acid in the other honey varietals may have hindered the probiotic’s capacity to adapt to the honey-yogurt food matrix compared to clover, but more studies are needed to further test microbial-organic acid interactions. Ultimately, the nutrient profile of clover honey may protect *B. animalis* through gastrointestinal digestion due to the presence of antioxidant phenolic compounds and reducing sugars, both of which can reduce the redox potential and protect anaerobic microbes, like *B. animalis*, from reactive oxygen species (28). Additionally, different honey varietals will have various antioxidative capacities, which can also be impacted by the gastric and intestinal environments (23). Future studies should test fractionated clover honey to identify the key fractions responsible for facilitating improved probiotic survivability.

A limitation of the honey nutritional composition profiles is that they only represent the undigested samples. More studies are needed to assess if these phenolic compounds have the same beneficial effect in isolation for *B. animalis* survivability after intestinal digestion compared to the other varietals. Another limitation was not performing antioxidant assays and measuring enzymatic activity of the honey varietals after each stage of digestion to assess any shifts from baseline in the antioxidative capacity. As noted in previous studies, the antioxidative capacity of honey is affected by the different environments presented during gastric and intestinal *in vitro* digestion (23). Therefore, future studies should track the bioactive activity of honey varietals as it goes through the different stages of digestion. Also, the commercial yogurt studied contained the probiotic, *B. animalis,* however, it also included the other live starter cultures that are standard in yogurt fermentation (i.e., *L. bulgaricus*, *L. lactis*, and *S. thermophilus*). Thus, there may have been cross-interactions between the probiotic and the starter culture during the baseline preparation, such as commensalism between microbes to break down certain components of the honey varietals that *B. animalis* could, perhaps, not do alone. As our work aimed to investigate the culinary pairing of yogurt and honey, this was beyond the scope of this work. However, additional work is necessary to understand how standard yogurt or yogurt with other probiotics may be impacted when paired with honey. Also, follow-up studies that aim to understand how *B. animalis* performs outside of the yogurt matrix would benefit by including another control sample that includes yogurt with only *B. animalis*, excluding the other yogurt cultures.

In summary, of the four honey varietals (alfalfa, buckwheat, clover, and orange), clover honey mixed with yogurt significantly increased *B. animalis* survivability after complete *in vitro* gastrointestinal digestion compared to three control treatments (yogurt with added sucrose, added water, and undiluted). Furthermore, clover honey tested at different dosages (0, 7, 10.5, 14, 17.5, 21, 28, 42 g per 170 g yogurt) demonstrated significant improvement in survivability for *B. animalis* using 21, 28, and 42 g per 170 g of yogurt compared to the control. This study demonstrated a ratio of two functional foods (honey:yogurt) supports *B. animalis’* survivability through *in vitro* digestion. High quality randomized controlled clinical trials are needed to determine if honey and probiotic honey pairing enhance probiotic survival and downstream benefits to the host.

## Abbreviations

TPTZ: 2,4,6-Tris(2-pyridyl)-s-triazine
DPPH: 2,2-Diphenyl-1-picrylhydrazyl
CFU: colony forming units
FRAP: Ferric-reducing antioxidant power
GAE: gallic acid equivalents
LAB: Lactic acid bacteria
AAE: L-ascorbic acid equivalent
MRS: De man Rogosa and Sharpe
TE: Trolox equivalents
TPC: total phenolic content

## Acknowledgments

We acknowledge Megan Hung, Noah Simon, and Muskaan Sawhney of the Nutrition and Human Microbiome Laboratory for their technical assistance with the assays and the *in vitro* digestion experiments. We also acknowledge Yan Zhu, Ronghua Liu, and Lili Mats of the Guelph Research and Development Centre, Agriculture & Agri-Food Canada for their contributions to the honey composition analyses.

## References

1. McKinley MC. The nutrition and health benefits of yoghurt. Int J Dairy Technol 2005;58:1–12.

2. Fazilah NF, Ariff AB, Khayat ME, Rios-Solis L, Halim M. Influence of probiotics, prebiotics, synbiotics and bioactive phytochemicals on the formulation of functional yogurt. J Funct Foods 2018;48:387–99.

3. Widyastuti Y. The role of lactic acid bacteria in whiskey brewing. Food Nutr Sci 2012;90:324–8.

4. Arena MP, Caggianiello G, Russo P, Albenzio M, Massa S, Fiocco D, Capozzi V, Spano G. Functional starters for functional yogurt. Foods 2015;4:15–33.

5. Chen C, Zhao S, Hao G, Yu H, Tian H, Zhao G. Role of lactic acid bacteria on the yogurt flavour: A review. Int J Food Prop [Internet] Taylor & Francis; 2017;20:S316–30. 10.1080/10942912.2017.1295988

6. Binda S, Hill C, Johansen E, Obis D, Pot B, Sanders ME, Tremblay A, Ouwehand AC. Criteria to Qualify Microorganisms as “Probiotic” in Foods and Dietary Supplements. Front Microbiol 2020;11:1–9.

7. Hill C, Guarner F, Reid G, Gibson GR, Merenstein DJ, Pot B, Morelli L, Canani RB, Flint HJ, Salminen S, et al. Expert consensus document: The international scientific association for probiotics and prebiotics consensus statement on the scope and appropriate use of the term probiotic. Nat Rev Gastroenterol Hepatol 2014;11:506–14.

8. Reid G, Gadir AA, Dhir R. Probiotics: Reiterating what they are and what they are not. Front Microbiol 2019;10:1–6.

9. Marco ML, Sanders ME, Gänzle M, Arrieta MC, Cotter PD, De Vuyst L, Hill C, Holzapfel W, Lebeer S, Merenstein D, et al. The International Scientific Association for Probiotics and Prebiotics (ISAPP) consensus statement on fermented foods. Nat Rev Gastroenterol Hepatol Springer US; 2021;18:196–208.

10. Meybodi NM, Mortazavian AM, Arab M, Nematollahi A. Probiotic viability in yoghurt: A review of influential factors. Int Dairy J 2020;109.

11. Yerlikaya O. Probiotic potential and biochemical and technological properties of Lactococcus lactis ssp. lactis strains isolated from raw milk and kefir grains. J Dairy Sci [Internet] American Dairy Science Association; 2019;102:124–34. Available from: 10.3168/jds.2018-14983

12. Quigley EMM. Chapter 13 - Bifidobacterium animalis spp. lactis A2 - Floch, Martin H [Internet]. The Microbiota in Gastrointestinal Pathophysiology. Elsevier Inc.; 2017. 127–130 p. Available from: https://www.sciencedirect.com/science/article/pii/B9780128040249000136

13. Hill C, Tancredi DJ, Cifelli CJ, Slavin JL, Gahche J, Marco ML, Hutkins R, Fulgoni VL, Merenstein D, Sanders ME. Positive Health Outcomes Associated with Live Microbe Intake from Foods, Including Fermented Foods, Assessed using the NHANES Database. J Nutr Elsevier BV; 2023;

14. Sanz ML, Polemis N, Morales V, Corzo N, Drakoularakou A, Gibson GR, Rastall RA. In vitro investigation into the potential prebiotic activity of honey oligosaccharides. J Agric Food Chem 2005;53:2914–21.

15. Chick H, Shin HS, Ustunol Z. Growth and acid production by lactic acid bacteria and bifidobacteria grown in skim milk containing honey. J Food Sci 2001;66:478–81.

16. Brodkorb A, Egger L, Alminger M, Alvito P, Assunção R, Ballance S, Bohn T, Bourlieu-Lacanal C, Boutrou R, Carrière F, et al. INFOGEST static in vitro simulation of gastrointestinal food digestion. Nat Protoc 2019;14:991–1014.

17. Li H, DZ, LR, LS, TR. Bioaccessibility, in vitro antioxidant activities and in vivo anti-inflammatory activities of a purple tomato (Solanum lycopersicum L.). Food Chem 2014;159:353–60.

18. Arany CB, Hackney CR, Duncan SE, Kator H, Webster J, Pierson M, Boling JW, Eigel WN. Improved recovery of stressed Bifidobacterium from water and frozen yogurt. J Food Prot 1995;58:1142–6.

19. Lapierre L, Undeland P, Cox LJ. Lithium Chloride-Sodium Propionate Agar for the Enumeration of Bifidobacteria in Fermented Dairy Products. J Dairy Sci [Internet] Elsevier; 1992;75:1192–6. Available from: 10.3168/jds.S0022-0302(92)77866-7

20. Vernazza CL, Gibson GR, Rastall RA. Carbohydrate preference, acid tolerance and bile tolerance in five strains of Bifidobacterium. J Appl Microbiol 2006;100:846–53.

21. Shin HS, Ustunol Z. Carbohydrate composition of honey from different floral sources and their influence on growth of selected intestinal bacteria: An in vitro comparison. Food Research International 2005;38:721–8.

22. White JW Jr, Riethof ML, Subers MH, Kushnir I. Composition of American honeys. US, Dep Agric, Tech Bull 1962;1–124.

23. Seraglio SKT, Valese AC, Daguer H, Bergamo G, Azevedo MS, Nehring P, Gonzaga LV, Fett R, Costa ACO. Effect of in vitro gastrointestinal digestion on the bioaccessibility of phenolic compounds, minerals, and antioxidant capacity of Mimosa scabrella Bentham honeydew honeys. Food Research International 2017;99:670–8.

24. Alaerjani WMA, Abu-Melha S, Alshareef RMH, Al-Farhan BS, Ghramh HA, Al-Shehri BMA, Bajaber MA, Khan KA, Alrooqi MM, Modawe GA, et al. Biochemical Reactions and Their Biological Contributions in Honey. Molecules. MDPI; 2022.

25. Lichtenberg-Kraag B. Evidence for correlation between invertase activity and sucrose content during the ripening process of honey. J Apic Res International Bee Research Association; 2014;53:364–73.

26. Szwajgier D, Jakubczyk A. Corresponding author-Adres do korespondencji: Dr inż Biotransformation of ferulic acid by lactobacillus acidophilus K1 and selected Bifidobacterium strains. ACTA Acta Sci Pol, Technol Aliment. 2010;9:45–59. Available from: www.food.actapol.net

27. Reverón I, de las Rivas B, Muñoz R, López de Felipe F. Genome-wide transcriptomic responses of a human isolate of Lactobacillus plantarum exposed to p-coumaric acid stress. Mol Nutr Food Res 2012;56:1848–59.

28. Favarin L, Laureano-Melo R, Luchese RH. Survival of free and microencapsulated Bifidobacterium: Effect of honey addition. J Microencapsul 2015;32:329–35.

